# Acceptability of active case finding with a seed-and-recruit model to improve tuberculosis case detection and linkage to treatment in Cambodia: a qualitative study

**DOI:** 10.1101/514208

**Authors:** Sovannary Tuot, Alvin Kuo Jing Teo, Danielle Cazabon, Say Sok, Mengieng Ung, Sangky Ly, Sok Chamreun Choub, Siyan Yi

## Abstract

**Background:** With support of the national tuberculosis (TB) program, KHANA has implemented an innovative approach using a seed-and-recruit model to actively find TB cases in the community. The intervention engaged community members including TB survivors to recruit people with TB symptoms in a ‘snowball’ approach for screening and linkage to treatment. This study aims to explore the acceptability of active case finding with the seed-and-recruit model in detecting new TB cases and determine the characteristics of successful seeds.

**Methods:** This qualitative study was conducted in four provinces (Banteay Meanchey, Kampong Chhnang, Siem Reap, and Takeo) in Cambodia in 2017. Fifty six in-depth interviews and ten focus group discussions were conducted to gain insights into the acceptability, strengths, and challenges in implementing the model. Transcripts were coded and content analyses were performed.

**Results:** The seed-and-recruit active case finding model was generally well-received by the study participants. They saw the benefits of engaging TB survivors and utilize their social network to find new TB cases in the community. The social embeddedness of the model within the local community was one of the major strengths. The success of the model also hinges on the integration with existing health facilities. Having extensive social network, being motivated, and having good knowledge about TB were important characteristics of successful seeds. Study participants reported challenges in motivating the recruits for screening, logistic capacities, and high workload during implementation. However, there was a general consensus that the model ought to be expanded.

**Conclusions:** These findings indicate that the seed-and-recruit model should be fine-tuned and scaled up as part of the national TB Program to increase TB new case detection in Cambodia. Further studies are needed to more comprehensively evaluate the impacts and cost-effectiveness of the model in Cambodia as well as in other resource-limited settings.

## Introduction

Cambodia is one of the 30 countries with a high burden of tuberculosis (TB) [1]. In 2016, the national incidence and prevalence rates of all forms of TB were 345/100,000 population and 668/100,000 population, respectively [2]. These rates have reduced substantially since 1990, and a similar decline was observed in the TB mortality rate [2,3]. Cambodia has made notable progress in the fight against TB by achieving a treatment success rate of 94%, one of the highest among the 30 high TB burden countries [4]. However, the successes are impeded by a significant proportion of under-diagnosed cases. Globally, it is estimated that 36% of TB cases were undiagnosed in 2017, and a similar proportion is observed in Cambodia [5–7].

In Cambodia, TB disproportionately affects people living with HIV, household and close contacts of bacteriologically confirmed cases, elderly above the age of 55, diabetics, prisoners, migrant workers, the poor, and those living in rural communities [2]. Despite a well-established national TB infrastructure, these key populations still face geographical, economic, social, and biomedical barriers to diagnosis and treatment [2,8]. In 2016, Ngin and colleagues reported that people with TB received inadequate consultation at both health centers and referral hospitals, and routinely experienced delays in diagnosis and treatment initiation [9]. Furthermore, health center staff, laboratory staff, and village health support groups were found to be insufficiently motivated in the conduct of their work [9].

Traditionally, TB cases are captured and passively notified when patients present themselves to a health facility. To further find undiagnosed cases and curb TB transmission, the National Center for Tuberculosis and Leprosy Control (CENAT) has adopted a proactive approach to find cases and promptly link them to care [5,7,10]. Despite the evidence of feasibility and cost-effectiveness of implementing active case finding among urban poor and rural settings in Cambodia [10,11], TB case finding remains a great challenge due to limited resources, geographical barriers, and social stigma. Finding new TB cases remains a great challenge given the stigmatization associated with TB and its highly contagious nature [12–14].

Empirical evidence suggests both efficiency and effectiveness in using a “snowball” approach to engage hard-to-reach key populations for HIV in many countries, including Cambodia [15–19]. However, little is known about the feasibility, acceptability, and effectiveness of this approach in improving the detection of new TB cases. Given the comparable hidden nature of HIV and TB populations, it is a notion worth exploring. To address this knowledge gap, we implemented active case finding with a seed-and-recruit model – a system of rapid, targeted low-cost community social mobilization, involving people who have themselves had TB (TB survivors), to increase case finding – in four national priority provinces in Cambodia. TB survivors were recruited as seeds to identify people with TB symptoms in their community. Newly diagnosed cases then became recruiters to recruit other people to be screened for TB in a snowball approach. This study aims to document the acceptability of the active case finding with the seed-and-recruit model in detecting new TB cases and determine the characteristics of successful seeds.

## Materials and Methods

### Ethics statement

This study was approved by the National Ethics Committee for Health Research (N. 238 NECHR) of the Ministry of Health in Cambodia. A letter of support was obtained from each provincial health department of the four provinces where the intervention was implemented. Verbal informed consent was requested from each participant before the data collection. Data collection was subsequently conducted in private places, and confidentiality was ensured by removing all personal identifiers from the final transcripts and field notes.

### Settings and study period

This qualitative study was conducted from November to December 2017 in four provinces where the seed-and-recruit model was implemented – Banteay Meanchey, Kampong Chhnang, Siem Reap, and Takeo.

### Sampling and participant recruitment

We conducted 56 in-depth interviews (IDIs) with lay counselors, TB program staff at health centers and referral hospitals, village health support groups, seeds, field staff, and TB patients. Also, ten homogenous focus group discussions (FGDs) were conducted with four groups of participants – TB patients, diabetics, elderly age 55 and above, and representatives from the general population (those aged between 15 to 54 years, including TB survivors). A stratified purposive sampling method was employed to recruit the study participants. Potential participants for the IDIs and FGDs were invited either in-person or via the telephone and email by the research team and lay counselors, respectively.

### Data collection

Data collection was performed by 10 data collectors who had extensive experience in qualitative research and were trained and closely supervised by the principal investigators. Information on the study and its objectives were provided verbally to potential participants. Interviews were arranged with those who agreed to partake at a time and location of convenience. The IDIs and FGDs took between 30 minutes to one hour and were audio-recorded. Participation was voluntary, and a token of appreciation (USD$5) were given to participants at the end of the interview.

The IDIs and FGDs were conducted using a semi-structured guide in Khmer. The guide comprised of broad themes to understand the process, feasibility, acceptability, and strengths of the model in TB case detection and linkage to care. We also sought to understand the challenges faced by seeds in implementing this intervention and in identifying the characteristics of seeds that may potentially contribute to the success of this intervention. The guide was pilot-tested before implementation. Individuals who participated in the pilot test were excluded from the main study.

### Data management and analyses

All data were transcribed verbatim into Khmer by members of the research team. Personal information was removed from the final transcripts to ensure anonymity and confidentiality. Transcripts were coded by a research team member in NVIVO 11 (QSR International). The initial themes based on the semi-structured guide were used to develop a codebook. We also performed content analyses to identify emerging categories, themes, and common and divergent patterns pertinent to the objectives of the study. We described socio-demographic information of the study participants using STATA 14 (StataCorp, LP, Texas, USA).

## Results

### Characteristics of study participants

In total, 120 respondents participated in the study. We conducted 56 IDIs and 64 individuals representing four key population groups in 10 homogenous FGDs – elderly above the age of 55 (four FGDs), diabetics (one FGD), people living with HIV (one FGD), and the general population (e.g., people below the age of 55, TB survivors) (four FGDs). Of the 56 IDIs, more than half of the participants were male. The median age was 51 (range 17-88). One-third of the FGD participants were male, and the median age was 54 years (range 17-78). Characteristics of the participants in IDIs and FGDs are summarized in Table 1.

**Table 1.**
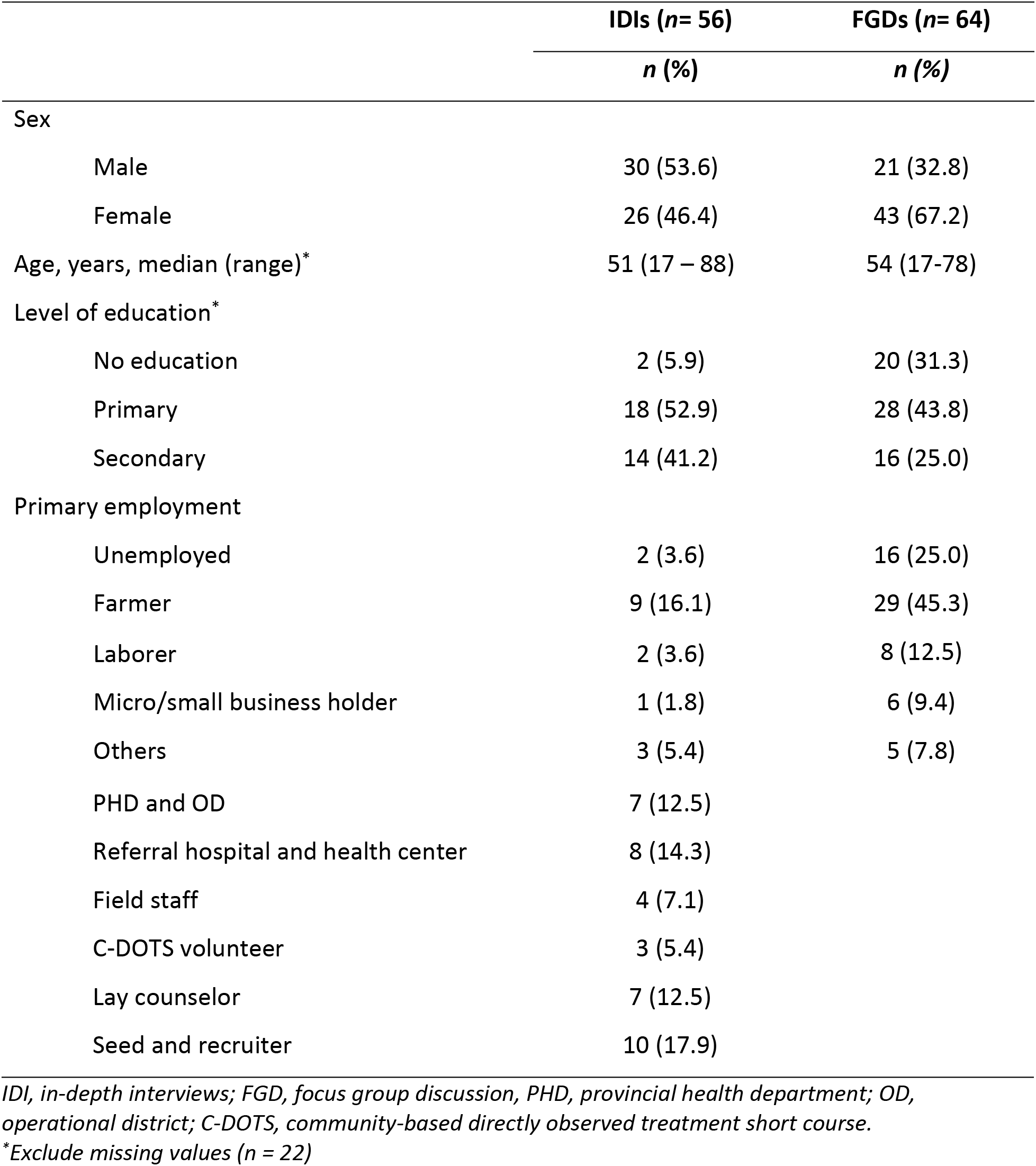
Characteristics of participants in in-depth interviews and focus group discussions.

|  | IDIs (*n*= 56) | FGDs (*n*= 64) |
| --- | --- | --- |
|  | *n* (%) | *n* (%) |
| Sex |  |  |
| Male | 30 (53.6) | 21 (32.8) |
| Female | 26 (46.4) | 43 (67.2) |
| Age, years, median (range)* | 51 (17 – 88) | 54 (17-78) |
| Level of education* |  |  |
| No education | 2 (5.9) | 20 (31.3) |
| Primary | 18 (52.9) | 28 (43.8) |
| Secondary | 14 (41.2) | 16 (25.0) |
| Primary employment |  |  |
| Unemployed | 2 (3.6) | 16 (25.0) |
| Farmer | 9 (16.1) | 29 (45.3) |
| Laborer | 2 (3.6) | 8 (12.5) |
| Micro/small business holder | 1 (1.8) | 6 (9.4) |
| Others | 3 (5.4) | 5 (7.8) |

|  |  |
| --- | --- |
| PHD and OD | 7 (12.5) |
| Referral hospital and health center | 8 (14.3) |
| Field staff | 4 (7.1) |
| C-DOTS volunteer | 3 (5.4) |
| Lay counselor | 7 (12.5) |
| Seed and recruiter | 10 (17.9) |

*IDI, in-depth interviews; FGD, focus group discussion, PHD, provincial health department; OD, operational district; C-DOTS, community-based directly observed treatment short course.*
*\*Exclude missing values (n = 22)*

**Table 2.**
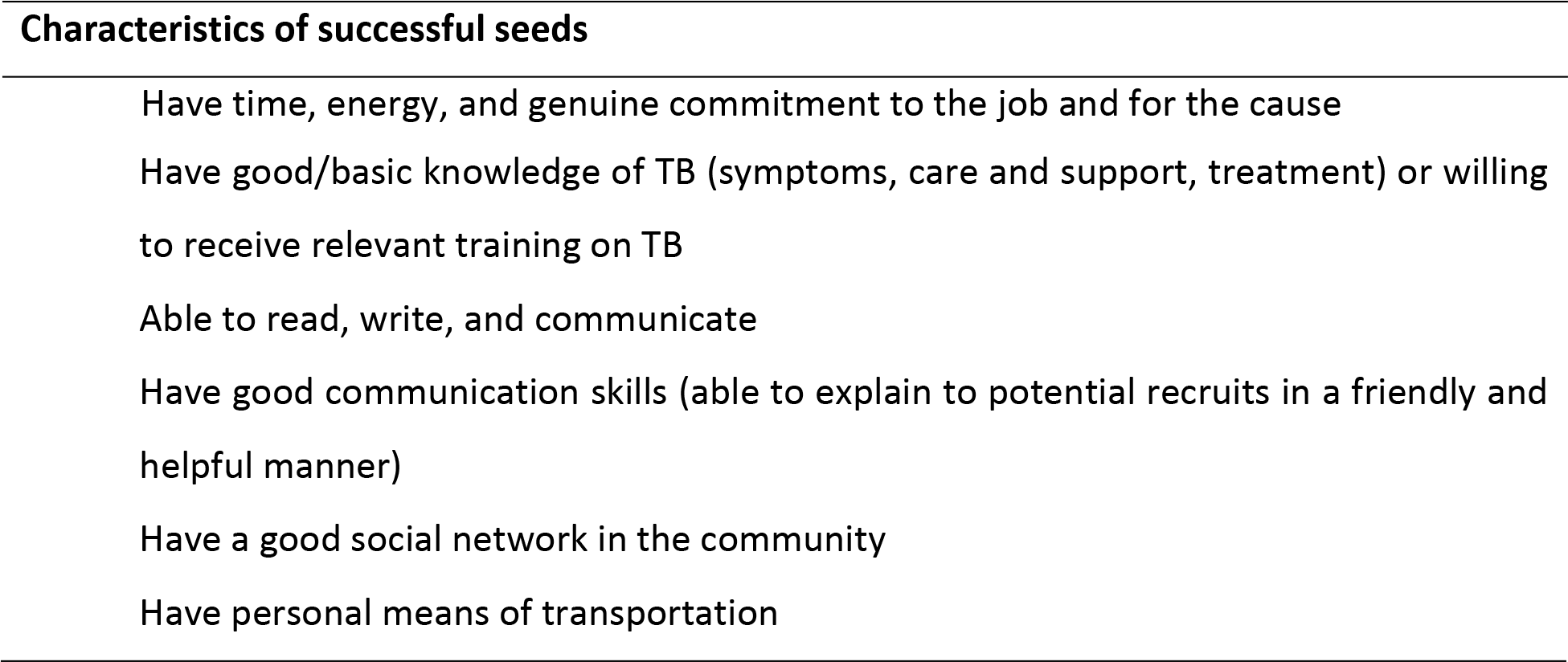
Characteristics of successful seeds in recruiting people at higher risks and producing high-yield TB screening.

### Acceptability of the seed-and-recruit model

In general, study participants found the seed-and-recruit model helpful for both the patients and the community. They appreciated that people who are sick would be screened and treated on time through this model. Hence, they perceived that the model must be expanded to halt transmission and eliminate TB from the community.

> “We do not want to transmit TB to people in our families and neighborhoods, and we want to eliminate TB from our community.” (IDI with elderly from Siem Reap)

The seed-and-recruit model mobilizes communities to seek people with TB symptoms and empowers them to be conscious about their health.

> “I think the model is good for me. Sometimes we forget to take care of our health, and they are a good reminder for us to check up our health.” (IDI with people living with HIV from Banteay Meanchey)

The integration of the model into the existing public health care systems was well-received by the study participants. The health centers were well-maintained and located conveniently close to their residences. They also commended the staff for their professionalism.

> “Health center is generally convenient and close to the place of residence of the patients.” (IDI with health center staff from Banteay Meanchey)

All key stakeholders agreed that having adequate TB screening facilities and health personnel at the health facilities are crucial for the success of the model. Generally, there are no shortage of staff and TB screening facilities at most locations.

> “Personnel and facilities do not seem to be an issue because the number of people seeking screening is still relatively low.” (IDI with an operational district staff from Kampong Chhnang)

The project staff, lay counselors, seeds, and health care providers saw a lot of benefits of engaging TB survivors (and the snowball approach, more broadly) to recruit potential people with TB. The seeds could easily identify people with TB symptoms for screening and encouraged them to seek care.

> “It is acceptable as the networks and seeds are so active in looking for the potential recruits.” (IDI with a community-based DOTS staff from Banteay Meanchey)

### Strengths of the model

Virtually, all respondents saw strengths in the model. Study participants expressed that the model encouraged early diagnosis and treatment. Patients were also duly guided during the screening and treatment process by the seeds at no cost.

> “A positive thing (about the project) is that our villagers are alerted earlier before TB gets worse. It is good that they come to search for us and guide us through the screening and treatment and to educate us too. We do not need to pay a cent.” (FGD with an elderly from Banteay Meanchey).

The ability of the model to identify more TB cases through trained seeds and recruiters were identified as one of the main strengths of the model.

> ‘Mobilizing well-trained seeds (and recruiters), who can identify more TB cases and send them to take the test at a health facility, is one of the strengths of the model.” (IDI with a VHSG from Banteay Meanchey)

This peer-driven model was a ground-up and community-based intervention. Study participants expressed that this model played a pivotal role in behavior change through skills building, peer influence, motivation, and knowledge sharing. An intervention as such, nested in the community, was perceived to be sustainable.

> ‘Social embeddedness of the seeds and recruiters in the local community and hence the ability to share knowledge and experience with the potential recruits, leading to sustainability of the intervention.’ (IDI with a health center staff member from Takeo)

### Challenges in implementing the seed-and-recruit model

Nevertheless, difficulties in motivating potential patients to seek screening and treatment services was expressed as one of the major challenges in the model implementation. It was attributed to three key issues – personal commitments (work, household chores), denial, and misconception regarding TB. Furthermore, additional trips to the referral hospital for further testing were not well-received by potential patients due to the distance between the referral hospital and their residence.

> “Some people are very difficult to talk to. Some people are ignorant and stubborn. People aged 70 and above are hard to talk to because they think that no one in their families has ever had TB.” (IDI with a seed from Banteay Meanchey)

> “Lack of cooperation from potential recruits in further recruitment (e.g. because of old age, busyness with making a living or tending grandchildren) is the main challenge in the model implementation.” (IDI with a project staff member from Banteay Meanchey)

Some lay counselors and seeds reflected that the case-by-case incentivization scheme is not enticing for seeds/recruiters to perform to their full potential.

> “Why don’t they provide us with regular incentives? Why are we provided with incentives only when we can find a case? When I travel to educate other people, I also need to spend time on the work.” (IDI with a community DOTS volunteer from Siem Reap)

Higher up the echelon, some local project staff indicated the difficulty in recruiting and maintaining lay counselors due to workload and the lack of incentives. Some study participants expressed that it was challenging for one lay counselor to manage patients in two health centers well and evenly. There was a tendency for a lay counselor to oversee the health centers in his/her residential district (or even commune/village) more closely and neglect the other one.

> “When we first started, we found it hard to recruit staff, i.e., lay counselors, and the incentives are quite little too.” (IDI with an lay counselor from Kampong Chhnang)

There were also instances of limitations in the screening capacities in some health centers. Some participants stated that the limited number of staff and diagnostic infrastructures at the health facilities occasionally prevented timely management and process of samples, resulting in a delay of diagnosis. In the matter of consumables outage, study participants concurred that they generally did not face major issues. If shortages were to occur, communications with other partners were swift to resolve the problem.

> There is only one GeneXpert machine at the health facility, and it is used to conduct sputum test from our project as well as from another NGO and the health facility itself. Therefore, sometimes they cannot handle all of them on time. In one day, at most, it can handle 16-20 bottles. Sometimes we need to wait for half a month to get the result.” (IDI with a lay counselor from Siem Reap province)

> “Like ink for the GeneXpert machine, occasionally they (health facilities) may run of stock. However, we contacted the OD straight away, and it was immediately resolved.” (IDI with a local project officer from Banteay Meanchey)

### Nationwide scalability of the model

Study participants expressed that the model can be scaled up and expanded nationwide due to the potential impact on TB case detection and cost-effectiveness of the intervention. The project also met the national program’s mission to control TB in the country.

> “I think it can be scaled up because the model is at low cost, and when we search for the TB patients everywhere, the intervention can be effective and TB burden will be reduced.” (IDI with a community DOTS volunteer from Siem Reap)

> “The special feature of this model is that our seeds are TB survivors, and hence they know the symptoms well and what to talk to potential recruits. This approach can produce high yields with little budget” (IDI with a health center staff member from Takeo)

> “It is scalable to the national level. If it is properly executed, we are able to find many cases. This also aligns with the national guidance/principles, which want health centers to look for new cases” (IDI with a referral hospital staff member from Siem Reap)

Nonetheless, a concern about health facilities’ inability to handle the volume of workload if the intervention is scaled up was raised and a systematic solution ought to be explored.

> “In the future, when we can find more cases, the challenge could be that we could not handle the cases as we have limited number of staff. Maybe in that case, we will need to have a proper appointment system with the recruits.” (IDI with a health center staff member from Kampong Chhnang)

### Characteristics of successful seeds

The ability of good seeds in finding new TB cases and mobilizing potential recruits to take the screening test varied from seed to seed. Nevertheless, their achievements in mobilizing testing and yielding positive results were encouraging. Successful seeds/recruiters were driven by both intrinsic and extrinsic motivation. Many lay counselors, seeds, and recruiters were motivated by their wish to eliminate TB from their communities, terminate future transmission, and be socially responsible. Incentives, new skills, and knowledge acquisition were also mentioned as reasons to get involved in the project.

> “I want to eliminate the transmission of TB to other people, including children. I love this work because I used to see the suffering of my mother who went through a hard time when she was seriously ill (i.e., her lung was destroyed), and I do not want this to happen to other people.” (IDI with a seed from Siem Reap province)

In summary, we inferred the following characteristics of successful seeds from the study participants that could be emulated in implementing the model in the future.

## Discussion

Overall, the seed-and-recruit model that links community people to TB screening and treatment is deemed to be feasible and acceptable by the stakeholders consulted in this study. The appropriateness of applying the seed-and-recruit model in TB case detection and linkage to treatment is similar to the findings in studies that examine its application in HIV programs in Cambodia and elsewhere. In a study by Pitter and colleagues, a peer-driven intervention (PDI) to reach key populations for HIV (similar to the seed-and-recruit model) was found to be more effective (yield of HIV positive cases) compared to the same time period in the previous year, and it was positively received by the target populations and program implementers [20]. The study also concluded that the cost per HIV case detected in the intervention arm was three times lower than that of the comparator approach (outreach health counseling and treatment) [20]. In the United States, China, and Vietnam, PDI was found to reach a larger and more diverse pool of people who use drugs and significantly reduced HIV-associated risk behaviors among them [21,22]. Another study by Yi and colleagues on the use of the PDI program to recruit new HIV cases among transgender women also demonstrated the successful application of the approach in Cambodia [19,23].

The effectiveness of community-driven interventions to improve TB diagnosis and care has also been demonstrated. In Myanmar, community intervention was shown to increase case notification rates compared to household and neighborhood interventions [24]. The involvement of community volunteers in TB control was also found to increase the yield of diagnosis by 12% and reduce diagnostic delay in India and Nepal, respectively [25–27]. A patient-led active case finding strategy implemented in the Democratic Republic of Congo (DRC) reported an increase in TB diagnosis, and the intervention was highly acceptable by the communities, health facilities, and patient leaders. In a separate study in the DRC, peer education intervention was found to increase TB case notification rates by two-folds compared to controls, despite being conducted in conflict-affected settings [28]. Among people living with HIV in Nepal, peer-led TB case finding intervention resulted in a high yield of TB diagnoses [29]. Besides improvement in TB case notification rates, the peer-led intervention was also shown to improve knowledge on TB and prevention in a study in China [30]. Regarding acceptability of the intervention, respondents found a peer-led intervention to be beneficial and adequate and suggested future expansion[30], concurring with the findings of our study.

An operational research study on the PDI project in HIV conducted by KHANA highlighted challenges in the program implementation such as seed/recruiter drop-out, hard-to-reach key populations, the unattractiveness of the approach (or incentives) to the seeds and recruiters, loss to follow-up, and time constraints for key populations to visit health facilities [31]. While some of these challenges may be specific to the HIV program, our study found similar constraints, and these ought to be critically addressed through education, communication, and sustainable incentivization scheme should the intervention be scaled up nationwide. Efforts were also made to identify the characteristics of good seeds. In the scale-up phase, these findings would be incorporated into the training materials of project staff to identify good seeds efficiently. Coupled with structured and continuous training and monitoring, networks of competent seeds will be instrumental to the success of the project in the future.

The strengths of this study include the inclusion of multiple viewpoints sought from a wide range of high-value subjects – project and health facilities staff, patients, and key populations for TB – on the acceptability, feasibility, and challenges in the model implementation. These opinions and perspectives will contribute to the improvement of the intervention for future scale-up. To our knowledge, this is the first project that applied a seed-and-recruit model to TB case finding in Cambodia, and therefore embodies potential contribution to the pool of evidence-based interventions to TB case finding. Utilization of semi-structured interview guides for IDIs and FGDs ensured consistency and only one coder was involved in the analysis of qualitative transcripts, thus allowing a detailed understanding of the data. Lastly, data saturation for all topics was achieved. The findings of this study, however, must be considered in the context of several limitations. Perspectives of policymakers at the national level were not elicited. Hence, the discussion on nationwide scalability might not be comprehensive. Second, the applicability of the qualitative findings is limited to the health centers included in the pilot phase. Nevertheless, we believe that standpoints that we have collected and synthesized in this study are pivotal to improve the intervention for future scale-up.

Based on the findings from this evaluation, we recommend the following improvements to enhance the intervention for nationwide expansion. Firstly, as many of people with TB symptoms are affected by multiple complicities, which prevent them from seeking care and treatment, other health and social services should be integrated into the intervention. Provision of subsidy support during TB treatment, knowledge on diabetes prevention and care, old-age complication and care, and TB-related information dissemination may synergistically improve TB outcomes, as well as overall health and well-being of the population. Secondly, future programs should provide more logistical support (e.g., stationeries and consumables) for lay counselors, seeds, and recruiters to perform their duties. Support and encouragement in the form of supervision and opportunities to troubleshoot are required to ensure that staff personal health and wellbeing is taken care, and project implementation is on-track. A more comprehensive training program that includes TB knowledge, interpersonal, administrative, and management skills must be planned for the project to succeed. Furthermore, there is a need to devise a sustainable financial support scheme to recruit and motivate lay counselors, seeds, and recruiters. Thirdly, there is a need for the seed-and-recruit programming to strengthen engagement with relevant stakeholders at all levels to ensure that they are well-informed regarding the project. Their involvement is crucial to scale-up and sustain this intervention in the long run. Finally, there is also a need to devise a support plan and sustainable financial support scheme for health center and referral hospital staff for their services in this project.

## Conclusions

The seed-and-recruits model is well accepted by all stakeholders. The success of this model are anchored to the participatory nature of its design and implementation, the bottom-up driven community empowerment approach, and strong commitment from key stakeholders. We recommend that the scale-up of this intervention integrates other health and social services to the model; continues to support, train and incentivize lay counselors, seeds, and recruiters; and strengthens engagement with stakeholders at all levels. It is imperative that the project is fine-tuned and subsequently scaled-up countrywide to complement the national efforts to end TB by 2035.

## Acknowledgments

The authors wish to thank the participants in this study. We also thank the provincial health department, operational health districts, and health center staff for supporting this study.

## Supporting information

S1 File. Guide for In-depth Interviews and Focus Group Discussions

S2 File. COREQ checklist for interviews and focus groups

